# The structure of anticorrelated networks in the human brain

**DOI:** 10.1101/2022.05.10.491394

**Authors:** Endika Martínez Gutiérrez, Antonio Jiménez Marín, Sebastiano Stramaglia, Jesus M. Cortes

## Abstract

During the performance of a specific task or at rest, the activity of different brain regions shares statistical dependencies that reflect functional connections. While these relationships have been studied intensely for positively correlated networks, considerably less attention has been paid to negatively correlated networks, a.k.a. anticorrelated networks (ACNs). Here, we have addressed this issue by making use of two neuroimaging datasets: one of N=192 young healthy adults; and another of N=40 subjects that was divided into two groups of young and old participants. We first provided a full description of the anatomical composition of the different ACNs, each of which participated in distinct resting-state networks (RSNs). In terms of their frequency of participation, from highest to the lowest, the major anticorrelated brain areas are the precuneus, the anterior supramarginal gyrus and the central opercular cortex. Subsequently, by evaluating the more detailed structure of ACNs, we show it is possible to find significant differences in these in association with specific conditions, in particular by comparing groups of young and old participants. Our main finding is that of increased anticorrelation for cerebellar interactions in older subjects. Overall, our results give special relevance to ACNs and they suggest they may serve to disentangle unknown alterations in certain conditions, as might occur in neurodegenerative diseases at early onset or in some psychiatric conditions.

## Introduction

It is well known that anatomical networks, such as those built from diffusion tensor imaging (DTI), introduce statistical dependencies in the dynamics of connected regions, increasing their functional connectivity (FC)^1,2^. However, FC can also occur between regions with no direct anatomical connections^3,4^, due to the effects of common inputs or physiological elements, or the so-called indirect effects that refer to correlations between two regions that arise from neighboring regions^1,2,5–9^. Functionally connected networks can be decomposed into positively correlated networks and negatively correlated networks, the latter widely known as anticorrelated networks (ACNs). When derived from magnetic resonance imaging (MRI), some preprocessing steps might enhance the presence of ACNs^10–12^, while it has also been proposed that ACNs could fulfil a physiological role independently on the details of the preprocessing^13–17^. The most famous ACN is the default mode network (DMN), widely shown to be anticorrelated with multiple-task correlated networks^16,18^. ACNs might also facilitate task-switching, as shown for the Dorsal Attention Network (DAN)^19^, Salience Network (SN) and the Executive Control Network (ECN) when switching from the resting state to task performance^20,21^. Here, we extend these results by studying each resting state network (RSN) and its corresponding ACN, hypothesising that novel relationships can be found that differentiate two study groups by comparing the structure of the ACN, over and above the classic differences found when comparing them with positive correlated networks. To confirm our hypothesis, we use the example of aging and we assessed the differences between groups of old and young subjects in the ACN.

## Materials and Methods

### Participants

Data were selected from the 1,000 functional connectomes project and downloaded from the FCP Classic Data Table (available at http://fcon_1000.projects.nitrc.org/fcpClassic/FcpTable.html)^22^. To identify and analyse ACNs, we chose the *Beijing Eyes-Open Eyes-Closed II (Beijing EOEC2)* dataset^23^, selecting N=192 subjects with no history of neurological or psychiatric disorders (age range 18-26 years, mean 21.17, std. dev 1.83; males = 74). To analyse the differences in ACNs produced by aging, we chose The Max Planck Institute Leipzig Mind-Brain-Body dataset– LEMON^24,25^, selecting 20 young participants (range 20-25years; 10 male, 10 female) and 20 old participants (range 70-80 years; 10 male, 10 female).

### Brain imaging acquisition and analyses

#### Beijing dataset

The MRI data were acquired on a SIEMENS Trio 3-Tesla scanner at Beijing Normal University. Functional data were acquired using the following parameters: TR = 2s; volumes = 230; functional resolution was 3.125 × 3.125 × 3 mm with 64 × 64 × 36 voxels. The T1-weighted sagittal three-dimensional magnetization-prepared rapid gradient echo (MPRAGE) sequence was acquired using the following imaging parameters: 128 slices; TR = 2530 ms; TE = 3.39 ms; slice thickness = 1.33 mm; flip angle = 7°; inversion time = 1100 ms; FOV = 256 × 256 mm^2^.

#### LEMON dataset

MRI raw and pre-processed images are available at^26^ and although the full details of data acquisition are given at^25,27^, we summarize here the main acquisition parameters. High-resolution structural images were acquired through a 3D MP2RAGE sequence^28^ using the following parameters: voxel size = 1.0 mm isotropic, FOV = 256 × 240 × 176 mm, TR = 5000 ms, TE = 2.92 ms, TI1 = 700 ms, TI2 = 2500 ms, flip angle 1 = 4°, flip angle 2 = 5°, bandwidth = 240 Hz/Px, GRAPPA acceleration with iPAT factor 3 (32 reference lines), pre-scan normalization, duration = 8.22 min. Functional images were acquired through four rs-functional(f)MRI runs, all in an axial orientation using T2*-weighted gradient-echo echo planar imaging (GE-EPI) with multiband acceleration. The acquisition parameter for all four runs were: voxel size = 2.3 mm isotropic, FOV = 202 × 202 mm^2^, imaging matrix = 88 × 88, 64 slices with 2.3 mm thickness, TR = 1400 ms, TE = 39.4 ms, flip angle = 69°, echo spacing = 0.67 ms, bandwidth = 1776 Hz/Px, partial Fourier 7/8, no pre-scan normalization, multiband acceleration factor = 4,657 volumes, duration = 15 min 30 s. During the resting-state scans, the participants were instructed to remain awake with their eyes open and to fix their vision on a crosshair.

### Image preprocessing

Functional images of the two datasets were pre-processed using the Functional Connectivity (CONN v18b) toolbox^29^, applying the default pipeline for volume-based analyses that consists of: functional realignment and unwarping, translation to a common reference of functional images, slice timing correction, artefact detection tool based identification of outlier scans with a 97% percentile threshold, functional segmentation and normalization, translation to a common reference of structural images, structural segmentation and normalization, and functional smoothing with a Gaussian kernel of full width at half maximum equivalent to 8 mm. De-noising included regressing out 5 white matter components and 5 components of the cerebral spinal fluid, 12 realignment time series, scrubbing, linear de-trending and applying a band-pass filter between 0.008-0.09 Hz.

### Statistical analysis

We analysed eight different brain networks using seed-based correlation analysis (SBC). One seed was chosen within each of the networks available in the *networks* atlas incorporated into CONN^29^ and composing of 32 Regions of Interest (ROIs). In addition, a different CONN atlas was used for the anatomical description of significant clusters with 132 ROIs resulting from the combination of the FSL Harvard-Oxford atlas cortical & subcortical areas^30^ and the AAL cerebellar areas^31^. After applying a SBC, we built ACNs for each of the following RSNs: DMN, Frontoparietal Network (FPN), Sensorimotor Network (SMN), DAN, Visual Network (VN), Language Network (LN), SN, and Cerebellar Network (CN). The seeds used to generate each network were the medial Prefrontal Cortex (DMN), the lateral Prefrontal Cortex (FPN), the lateral Sensorimotor (SMN), the ipsilateral region (DAN), the medial Visual (VN), the Inferior Frontal Gyrus (LN), the Anterior Cingulate Cortex (SN), the Posterior region (CN), respectively.

A one-sample T-test was performed to analyse the ACNs and for group comparisons we performed two-sample T-tests. Voxel wise multiple correction was performed on the Beijing dataset using p-FDR-corrected (p-False Discovery Rate) 10^^-14^ and a peak to voxel p-FWE-corrected (p-Family Wise Error) 10^^-14^. Corrections were applied to the Lemon dataset using p-FDR-corrected 0.05 and cluster size p-FDR-corrected 0.05. The difference in the statistical threshold used in the two datasets was due to the differences in sample size between them.

## Results

Eight different RSNs were built using a SBC (figure 1), the: DMN, FPN, SMN, DAN, VN, LN, SN and CN.

**Figure 1.**
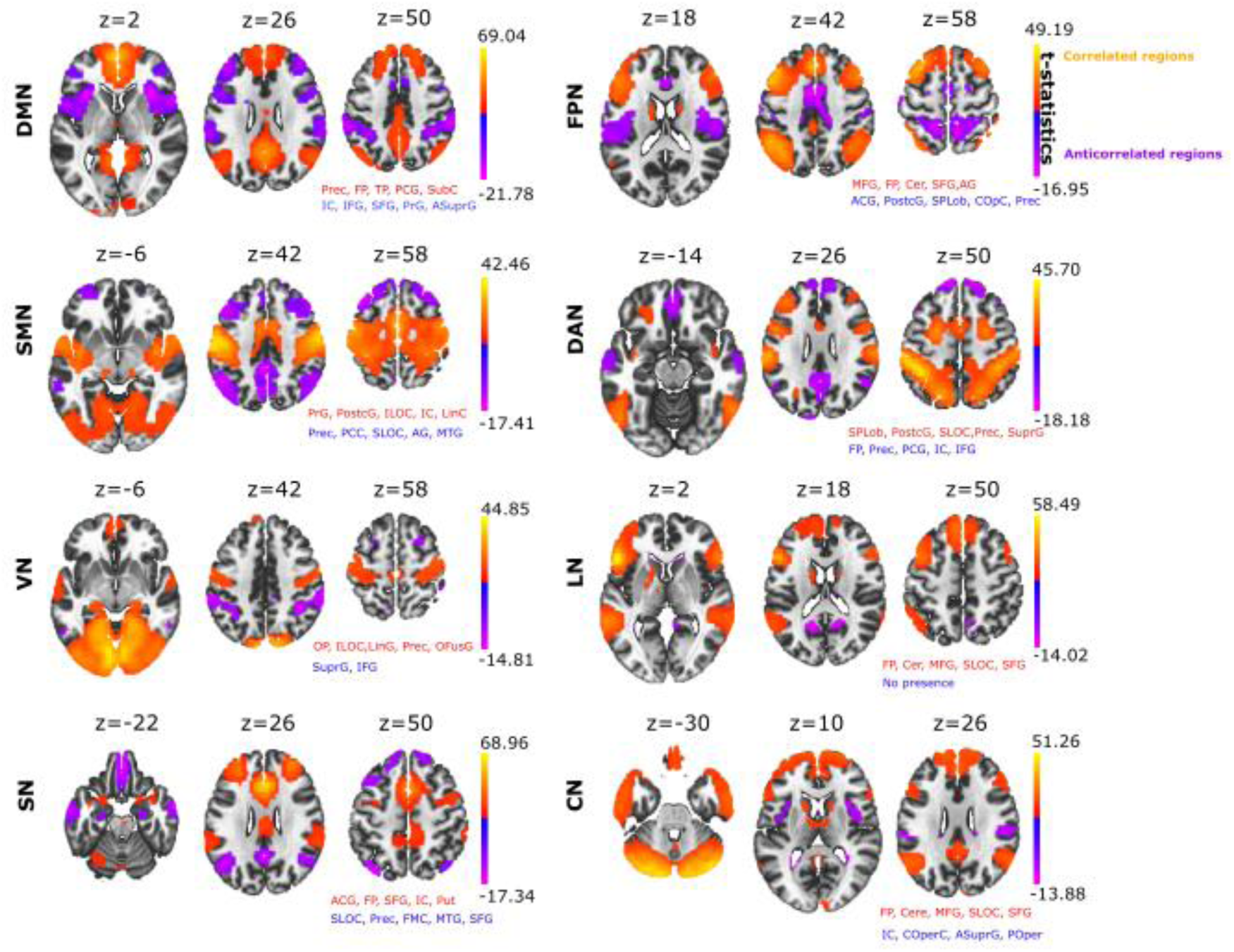
Seed based correlation analysis provides major resting state networks. Positive correlated networks (C>0; yellow-orange) and anticorrelated networks (C<0; blue-purple) for each of the eight networks: the Default Mode Network (DMN), the Fronto Parietal Network (FPN), the SensoriMotor Network (SMN), the Dorsal Attention Network (DAN), the Visual Network (VN), the Language Network (LN), the Salience Network (SN) and the Cerebellar Network (CN). For each of the networks, the text in red and blue indicates the names of the brain structures appearing in both the positive and negative correlated networks, respectively (the name of the structure appears if the brain map overlapped by 10% or more). Abbreviations: Precuneus (Prec), Frontal Pole (FP), Temporal Pole (TP), Posterior Cingulate Gyrus (PCG), Subcallosal Cortex (SubC), Insular Cortex (IC), Inferior Frontal Gyrus (IFG), Superior Frontal Gyrus (SFG), Precentral Gyrus (PrG), Anterior Supramarginal Gyrus (ASuprG), Middle Frontal Gyrus (MFG), Cerebellum (Cer), Anterior Cingulate Gyrus (ACG), Postcentral Gyrus (PostcG), Superior Parietal Lobule (SPLob), Central Opercular Cortex (COpC), Inferior Lateral Occipital Cortex (ILOC), Lingual Cortex (LinC), Superior Lateral Occipital Cortex (SLOC), Anterior and Posterior Supramarginal Gyrus (SuprG), Occipital Pole (OP), Lingual Gyrus (LinG), Occipital Fusiform Gyrus (OFG), Putamen (Put), Frontal Medial Cortex (FMC), Parietal Operculum (POper). Voxel wise multiple correction: p-FDR-corrected = 10^-14 and peak to voxel p-FWE corrected = 10^^-14^.

When we looked at which brain structures participated in all the different ACNs, we found that some structures were repeated more than others. We first determined the appearance of each structure across the entire set of ACNs. The anatomical structure that was most frequently present in the ACNs was the precuneus, which displayed significant anticorrelation in four of the eight RSNs analysed (Table 1), appearing most strongly in the SMN and reducing progressively through the DAN, SN and FPN. The next structure with the second most relevant role in ACNs was the Anterior Supramarginal Gyrus, which was also associated with four of the eights RSNs, namely the VN, DMN, FPN and CN. The subsequent dominant structure was the Central Opercular Cortex having a role in the ACNs associated with the DMN, FPN and CN.

**Table 1.**
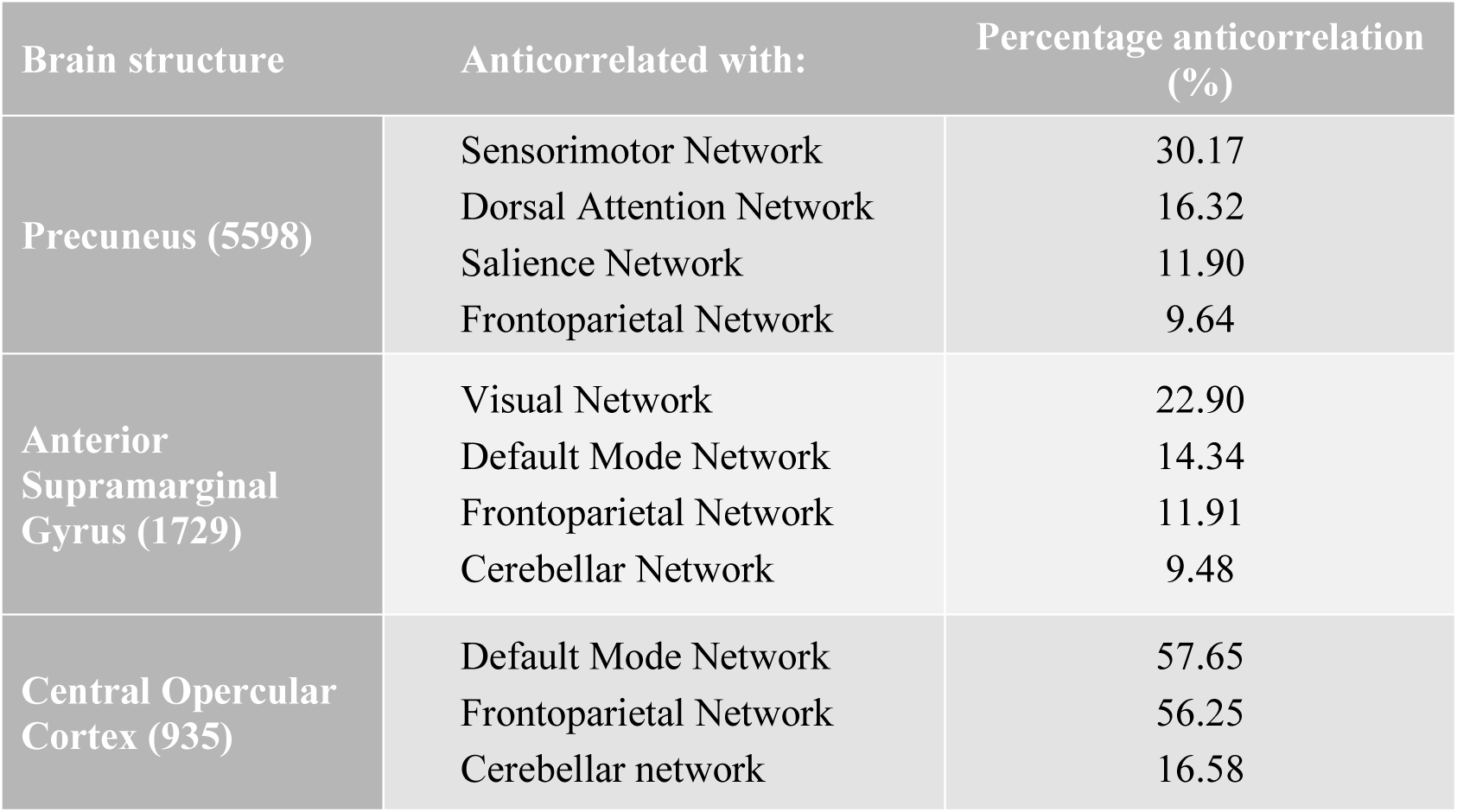
Major structures fulfilling an anticorrelated role in different RSNs. The Table shows the structures most frequently present in ACNs and their volumes (in brackets), measured in the number of voxels equal to 2× 2 × 2 mm3. The name of the RSN with which the structures establishes an anticorrelation, and the relative degree of overlap (%) between the brain structure and each RSN is also shown.

| Brain structure | Anticorrelated with: | Percentage anticorrelation (%) |
| --- | --- | --- |
| **Precuneus (5598)** | Sensorimotor Network | 30.17 |
|  | Dorsal Attention Network | 16.32 |
|  | Salience Network | 11.90 |
|  | Frontoparietal Network | 9.64 |
| **Anterior Supramarginal Gyrus (1729)** | Visual Network | 22.90 |
|  | Default Mode Network | 14.34 |
|  | Frontoparietal Network | 11.91 |
|  | Cerebellar Network | 9.48 |
| **Central Opercular Cortex (935)** | Default Mode Network | 57.65 |
|  | Frontoparietal Network | 56.25 |
|  | Cerebellar network | 16.58 |

In relation to our hypothesis that the structure of an ACN can differentiate between brain conditions, we asked whether ACNs could distinguish relevant aspects of aging. To achieve this goal, we analysed the ACNs for the eight different RSNs defined in a sample of young and old participants. Notably, we found that cerebellar regions showed greater anticorrelation with multiple brain areas in older adults than in younger adults in four of the eight ACNs analysed. Specifically, stronger cerebellar anticorrelations were evident in the older population in the SN, DAN, FPN and LN. We also found that in the DMN there was a greater anticorrelation with cerebellar regions in young adults (Figure 2), in contrast to what was observed in the rest of the network where the cerebellum had a stronger anticorrelated role in the older individuals. Moreover, we also found that the precentral gyrus had significantly stronger anticorrelations with the DAN in the young as opposed to the older participants, although the opposite contribution was found for the caudate and cerebellum, where they were more strongly anticorrelated with the DAN in older participants than in younger participants. When comparing the ACNs associated to the cerebellar network, we found that several brain structures participated more strongly ACN in the older participants, such as the precentral gyrus, frontal pole, putamen and supramarginal gyrus.

**Figure 2.**
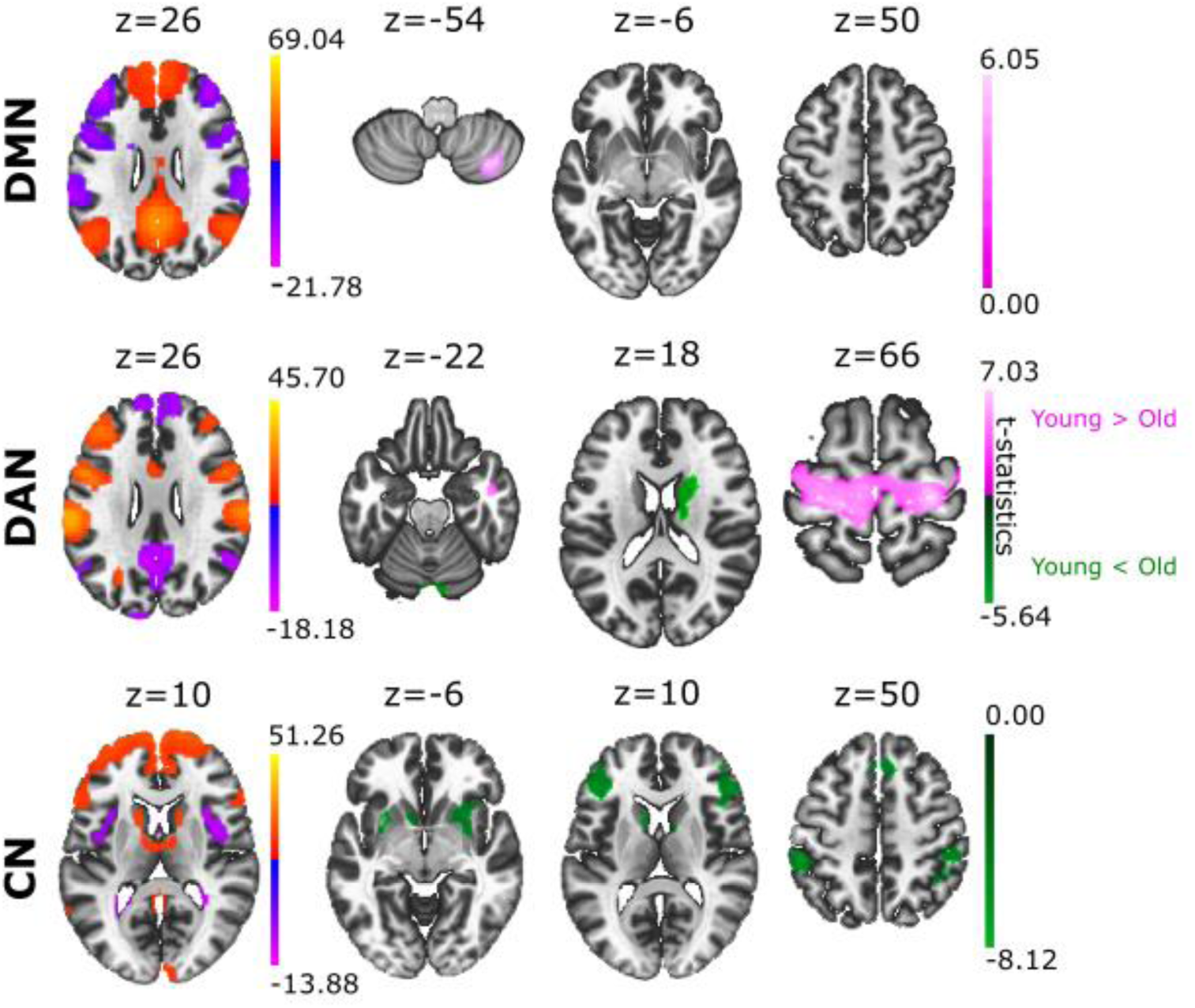
Group differences in the ACNs between young and old adults. In terms of the contrast Young > Old (coloured in pink), the different ACN maps provided significant differences in DMN and DAN. When comparing the groups with the contrast Young < Old (coloured in green), we found significant ACN differences in DAN and CN. Voxel wise multiple correction: p-FDR-corrected = 0.05 and cluster size p-FDR-corrected = 0.05.

## Discussion

Most studies of functional connectivity at rest have analysed positive correlation networks, meaning that ACNs have to some extent been neglected. Here, we describe the complete structure of ACNs, obtaining a different network for each of the well-known RSNs analysed. Our data show that the spatial distribution of anticorrelation structures is very heterogeneous, with the strongest contribution to anticorrelation found for the precuneus, the anterior supramarginal gyrus and the central opercular cortex.

The anticorrelating role of the precuneus as part of the DMN has been proposed previously, as DMN shows anticorrelation with the activation of multiple regions for tasks that demand attention and mental control^16,32–36^. The connectivity of the precuneus is extensive and widespread, involving cortical and subcortical structures that participate in the processing of highly integrated and associative information, rather than the direct processing of external stimuli^37^. Furthermore, the precuneus is functionally specialized for spatially guided behavioural processing^38^ and the activation of the precuneus precedes the onset of imagined movement, indicating that the precuneus may be involved in generating spatial information related to imagined peripheral and body movements^39^. Other studies have shown deactivation of the precuneus can be driven by different visual tasks, including visual attention^40^ and perception^41^, visual working memory and episodic memory^36^. This deactivation would be consistent with our findings, highlighting the critical role of the precuneus as the main anticorrelated hub across many human brain networks. The second structure with an important anticorrelation role was the anterior supramarginal gyrus, which overlaps with the SN. The supramarginal gyrus has been shown to play a key role in eliciting affective, self-other distinctions and empathic responses^42^. Anticorrelation between the SN and DMN has also been shown^43^, in part due to the fact that the SN is involved in externally directed attention demands for tasks that require cognitive control and where the insula reflects the main “core” for its implementation^21^. As part of the vestibular system, the supramarginal gyrus has also been seen to be involved in anticorrelation with the VN, integrating multisensory signals and playing an important role in visual integration^44^. In addition, we found anticorrelation for the supramarginal gyrus to the FPN (also known as the Central Executive Network), consistent with a Granger causality analysis that showed the SN mediates the switch between the DMN and FPN^45^. Our findings also reveal anticorrelation from the supramarginal gyrus to the CN, in agreement with previous studies relating the vestibular system and cerebellum^46^, with the cerebellum being crucial for the development of an internal model of action^47^, and the vestibular system relevant for perception, navigation and motor decision-making. Finally, the central opercular cortex is the third most relevant structure in terms of anticorrelation. Located in the Cingulo-Opercular Network (CON), it also represents a part of the SN^48^, and it is activated during the performance of some tasks like trial initiation and target detection, while it is also widely involved in a broad range of cognitive processes^21^.

In the second part of our work, we compared two different states in relation to the ACNs, being old with respect to being young. Previous studies showed a decrease in the intra-network functional connectivity and increased functional inter-network connectivity in older as opposed to younger adults^49,50^. Furthermore, decreased activation of the DMN has been seen in older adults^49^, as well as weaker anticorrelated interactions between the DMN and DAN in older as opposed to younger adults^51^. Our results show higher anti-correlation in the DMN of younger adults with the cerebellum lobule VII, which corresponds with cognitive demand functions^52^, and with cerebellum lobule VIII that is involved in sensorimotor tasks^6,52^. In relation to the cerebellar participation in the DAN, older adults also showed stronger anticorrelation in the cerebellum crus I, cerebellum crus II (corresponding to sensorimotor tasks) and the vermis VII that is involved in the proprioception of the body^53^. We also showed stronger caudate anticorrelations in the DAN for older adults, which is known to be implicated in motor processing^54^.

Overall, the findings of stronger anticorrelations to the CN in older adults might correspond to functional alterations that occur with age^55^, particularly in several cognitive functions like movement control, executive coordination or emotional regulation^6^ but also, it might be related to changes in cerebellar morphometric volume^56^. In the same way that multiple studies have correlated the activation of networks in task fMRI with those at rest for positively correlated networks^57–60^, future complementary studies should address the ACNs in task-specific fMRI, comparing them with those that we report here for the resting condition. In addition, our work provides a new method to study certain psychiatric or neurodegenerative conditions, comparing the structure of ACNs instead of positive correlated networks.

